# Advanced marker-assisted selection versus genomic selection in breeding programs

**DOI:** 10.1101/2023.02.20.529263

**Authors:** Bernd Degen, Niels Müller

## Abstract

Advances in DNA sequencing technologies allow the sequencing of whole genomes of thousands of individuals and provide several million single nucleotide polymorphisms (SNPs) per individual. These data combined with precise and high-throughput phenotyping enable genome-wide association studies (GWAS) and the identification of SNPs underlying traits with complex genetic architectures. The identified causal SNPs and estimated allelic effects could then be used for advanced marker-assisted selection (MAS) in breeding programs. But could such MAS compete with the broadly used genomic selection? This question is of particular interest for the lengthy tree breeding strategies. Here, with our new software SNPscan we simulated a simple tree breeding program and compared the impact of different selection criteria on genetic gain and inbreeding. Further, we assessed different genetic architectures and different levels of kinship among individuals of the breeding population. Interestingly, apart from progeny-testing, genomic selection (GS) using gBLUP performed best under almost all simulated scenarios. MAS based on GWAS results outperformed GS only if the allelic effects were estimated in large populations (ca. 10,000 individuals) of unrelated individuals. Notably, GWAS using 3,000 extreme phenotypes performed as good as the use of 10,000 phenotypes. Genomic selection increased inbreeding and thus reduced genetic diversity more strongly compared to progeny testing and GWAS-based selection. In conclusion, our analyses further support the potential of genomic selection for forest tree breeding and improvement although MAS may gain relevance with decreasing sequencing costs in the future.

## Introduction

Due to the advance of next generation sequencing technologies, the application of large marker sets with thousands of SNPs to estimate genomic breeding values (genomic prediction, genomic selection) has become a common practice in animal and plant breeding (Misztal et al. 2020, Sandhu et al. 2022). The effectiveness and accuracy of the genomic predictions have been studied with stochastic simulation models (Perez-Enciso et al. 2017) and deterministic models (Grattapaglia and Resende 2011). In contrast to the success of genomic selection, marker-assisted selection (MAS) could not fulfil the expectations in breeding programs (Grattapaglia and Kirst 2008, Kiszonas and Morris 2018) despite four decades of research and development (Nadeem et al. 2018). Initially, the main focus was on the search for quantitative trait loci (QTLs) defined as regions of the genome associated with a particular phenotypic trait. Most studies used linkage mapping in biparental populations to identify QTLs (Würschum 2012). In the beginning, QTL-studies were performed with a relatively small number (< 1000) of genetic markers (RAPDs, AFLPs, SSRs). The total amount of phenotypic variation explained by the QTLs was small due to the polygenic nature of most studied traits and due to the low proportion of loci segregating for the causal alleles in biparental populations. Further, the application of the QTLs to other unrelated material was a challenge.

With the advance of DNA sequencing techniques many more markers, especially SNPs, have been used to detect QTLs and the first genome-wide association studies (GWAS) in plants were done to detect causal SNPs in populations of unrelated individuals (Atwell et al. 2010). GWAS identified many more causal loci compared to QTL mapping (Alqudah et al. 2020). However, these studies were mostly done with arrays of only a few thousand SNPs (Alqudah et al. 2020), so again the SNPs identified in GWAS relied on linkage to the true causal variants and thus were difficult to use for unrelated individuals. Although the proportion of SNPs identified and the proportion of phenotypic variation explained increased a lot, it was still not enough to effectively estimate individual breeding values for a breeding program (Müller et al. 2017).

As demonstrated in human genetics, the use of whole-genome sequencing data can have a large impact on GWAS results (Wainschtein et al. 2022). Causal SNPs could be identified directly without relying on linkage or imputation and with increasing sample sizes more and more of the “missing heritability” could be recovered (Yengo et al. 2022). These results raise the question of whether an advanced MAS approach, based on these whole-genome GWAS, would perform equal or even better than genomic selection in a breeding program.

MAS and GS are of particular interest to tree breeding, because of the difficulty of phenotyping large and often heterogenous breeding populations and the long duration of breeding cycles. Even for the fast-growing Eucalyptus species a breeding cycle takes at least 8 years (Mphahlele et al. 2020) for most other tree species a breeding cycle takes decades. The limitation is caused by the late reproductive maturity and long phenotyping phase required for growth-related traits of at least one third of the rotation period. This is the reason why genomic selection has become so popular in tree breeding programs (Grattapaglia et al. 2018). MAS, however, could further facilitate the selection of seed donors in heterogenous forest stands that are unamenable to reliable phenotyping. To assess the potential of different breeding strategies, simulation studies can provide useful information (Liu et al. 2019). Here we used our new simulation program SNPscan to compare the expected performance of advanced MAS, genomic selection and the traditional selection criteria phenotype and progeny test in a simple tree breeding program. We studied the dynamics of genetic gain and inbreeding.

## Methods

### Simulation program SNPscan

Since decades, computer simulations are used to help breeders to make decisions on the choice of individuals for the next breeding cycle and on the different options for the breeding strategy (Bellmann and Ahrens 1966, Sun et al. 2011). Although, there are quite a few simulation tools and R-packages available to analyse different breeding strategies such as AlphaSimR (Gaynor et al. 2021), Xsim (Chen et al. 2022) and ADAM-Plant (Liu et al. 2019), there is still a gap for a user friendly windows program that enables simulations of simple genome-wide SNP data, implements user-defined genetic architectures and offers a broad set of selection criteria for forward simulations in breeding programs. Such functionalities are essential to optimise sample designs for the number of individuals and SNPs needed for accurate genomic predictions and marker-assisted selection based on GWAS results. Our program SNPscan aims to fill this gap. We use the program to compare different selection criteria in a simple tree breeding program and to gain insights into the effects of different possible breeding strategies.

With SNPscan the user defines distribution parameters for the generation of genomes and the genetic architecture of the trait. Then the program generates individual genomes and phenotypes according to these parameters. The option “Forward selection” offers various alternatives for the selection of parents for stochastic simulations of breeding cycles. The genomes of all individuals are stored in separate text files and can be aggregated and exported for further downstream analyses as files in phased HapMap-format.

#### Chromosomes and SNPs

SNPscan assumes diploid sets of chromosomes and co-sexual individuals (monoecious or hermaphroditic). The user defines the number of chromosomes, the total number of SNPs, the genome size and the average number of crossing-overs per chromosome. The SNPs are equally distributed over all chromosomes. The chromosomes have equal sizes. E. g., a genome with 20 chromosomes and a total of 2 million SNPs will have 100,000 SNPs per chromosome. The probability for a crossing-over is identical along the chromosomes. Other parameters to be specified by the user are the proportions of bi-allelic, tri-allelic and tetra-allelic SNPs as well as the distribution of frequencies of the common alleles at the SNPs. The default values are based on population re-sequencing data of different tree species, specifically beech (*Fagus sylvatica*), oak (*Quercus robur*) and ash (*Fraxinus excelsior*). For details on these genomic data see Sollars et al. (2017), Pfenninger et al. (2021), and Plomion et al. (2016).

#### Genetic architecture

The user defines the name of a trait, its mean phenotypic value in the founder or wild population, the variance of the phenotypes and the heritability of the trait. Then the number of causal SNPs is specified and one of three alternative functions for the distribution of the allelic effects selected: a) negative exponential distribution, b) normal distribution, or c) equal distribution.

#### Phenotypes

The phenotype of each individual *i* (*p*_i_) is computed as the sum of the mean phenotype of the population at the beginning (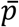 = parameter of the model), the genetic value (*g_i_*) and an environmental value (*e_i_*): 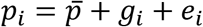

##### Genetic value (genomic breeding value)

The genetic value of individual *i* (*g_i_*) is calculated as the sum of all additive effects at each causal locus j (*a_ij_*) + a correction *m* to centralise the mean of all genetic values of the initial population to 0 multiplied with the scale factor *sf*: 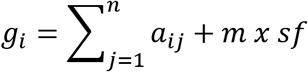

The scale factor (*sf*) is defined as: 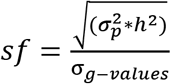

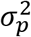= variance of the phenotypes in the population (parameter of the model)
*h^2^* = heritability (0-1, parameter of the model)
*σ_g-values_* = standard deviation of the genetic values of the population in the beginning

##### Environmental value

Environmental values (*e_i_*) are sampled from a normal distribution with a mean of 0 and the standard deviation of environmental effects: 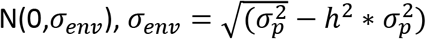

#### Forward selection

After the parents have been created, the user can run a forward selection by choosing among different options on how to select the parents for the next mating cycles:

“Phenotypes of adults”: Adults are ordered according to their phenotypes and the defined proportion of individuals are selected.

“Phenotypes of progenies”: For each adult a given number of offspring (“N progenies”) are simulated. The mating is random among all adults. The mean phenotype of the offspring is then used to rank the adults (estimate the genomic breeding values).

“GWAS estimates of allele effects”: This option is possible if a genome-wide association study (GWAS) has been performed on the simulation results and the estimated allelic effects stored. The program will ask for the according file. During the simulations, the breeding values of the adult individuals are estimated as the sum of the allelic effects at all identified associated SNPs.

“Genomic selection (gBLUP) single generation”: The program uses the function “kinship.BLUP” integrated in the R-package “rrBLUP” (Endelman 2011, R-Core-Team 2022). The algorithm computes “Predicted Genomic Breeding Values (PGBV)” for all individuals in the actual generation F with help of their phenotypes and the kinship-matrix of all individuals. The user can select the number of SNPs used for the predictions. As an option the causal SNPs can be excluded from these randomly selected SNPs. During the simulations SNPscan creates a subdirectory “R” in the project folder in order to store the input files, r-script for the calculation and the results.

“Genomic selection (gBLUP) cross generations”: As above but the phenotypes of all individuals of generation F-1 are used as training population and the individuals of the generation F are the test population. The algorithm computes “Predicted Genomic Breeding Values (PGBV)” for all individuals in generation F with help of phenotypes of generation F-1 and the kinship-matrix of all individuals in both generations.

#### Genome-wide association studies (GWAS)

SNPscan simulates parents and offspring that can be used for genome-wide association studies (GWAS). For this, simulated genomes of parents or offspring are stored as a HapMap-file and their phenotypes are stored in a text file. There are many different programs and R-scripts available to run GWAS. For the interaction with SNPscan we have selected the program Tassel Version 5.0 (Bradbury et al. 2007). The Tassel results on GWAS using the GLM and MLM algorithm can be loaded into SNPscan for post-ex analysis. For user defined thresholds of the association probabilities, the proportion of true and false positive associated SNPs are analysed. The estimated allelic effects are compared to the true effects using the Pearson’s correlation coefficient. Further correlation coefficients between true and estimated individual breeding values are computed considering the identified SNPs and their allelic effects. Finally, the user can store the allelic effects of all SNPs above the probability threshold as a file used for marker-assisted selection (MAS) in forward simulations with SNPscan.

#### Genetic diversity

Using 100 “dummy” loci with unique alleles for all individuals in the initial founder generation SNPscan computes important population genetic parameters:

“Inbreeding F (0-1)”: This is the inbreeding coefficient *F* defined as the probability that the two alleles of a homozygous genotype are identical by decent.
“Kinship (0-1)”: This is defined as the average probability that alleles at a locus of pairs of individuals are identical by decent.
“Rep.pop.size (*Np*)”: The reproductive effective population size is the effective number of parents contributing to the actual generation of offspring weighted by the relative fitness:
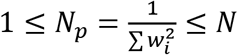 with (*w_i_*) = proportion of successful male and female gametes of each individual.
“Inbreed. Pop. size (Ne)”: The inbreeding effective population size: 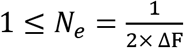, with ΔF = difference of inbreeding coefficient of current and last generation (Falconer and Mackay 1996).
“Ancestor diversity (*A_v_*)”: This is the effective number of genetically unrelated ancestors contributing to the actual generation of offspring. During the generation of the parent genomes, each individual is assigned unique alleles at 100 loci *i*. Thus, in the beginning of the simulation the diversity at these “ancestor alleles” is equal to the initial number of parents (N). *A_v_* is computed as the average diversity at all 100 ancestor loci:

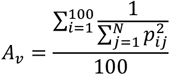

#### Scenarios

Here, we simulated a simple breeding program typical for a tree species with closed recurrent selection and separated generations of a monoecious or hermaphroditic, diploid species. The species had ten chromosomes and a genome of 500 million base pairs (Mbp). In all scenarios we simulated one million SNPs which have been equally distributed over the genome resulting in 100,000 SNPs per chromosome. We assumed a single target trait. The narrow sense heritability (h^2^) was 0.5. In each generation 1,000 trees were generated. In scenario 1 we had no burn-in. In scenario 2 and 3 we ran 20 generations as a burn-in with 100% of the males contributing to the random mating but only 10% randomly selected trees used for the harvest of seeds for the next generation. This burn-in created a gradually varying level of kinship among the individuals with an average of 0.04.

In scenario 1 and 2 the trait was encoded by 200 causal SNPs and in scenario 3 by 20 SNPs. The effects at these SNPs followed a standard normal distribution with a mean of 0 and a standard deviation of 1. There was random mating among all selected parents. The 10% top parents for the next breeding cycle were selected according to the following criteria (sub scenarios):

a. Phenotype of adults => *Phenotypes*
b. Breeding values estimated by progeny tests (100 offspring per tree generated by random mating among all 1000 trees) => *Progenies*
c. Single generation genomic selection with gBLUP using 10,000 randomly selected SNPs (excluding causal SNPs) recalculated in <u>each generation</u>. Here all individuals provided genotypes and phenotypes to the analysis and thus there was no separation between a training and a test data set. => *gBLUP-singleG*
d. Cross generations genomic selection with gBLUP using 10,000 randomly selected SNPs (excluding causal SNPs) recalculated for each generation with phenotypes of _Fn-1_ and genotypes of F_n-1_ and F_n_. Here the kinship-matrix and the phenotypes of F_n-1_ serves as training set and the <u>next generation</u> F_n_ is the test data set => *gBLUP-crossG*
e. Marker-assisted selection using estimated allelic effects of the 100 SNPs with the lowest p-values in a genome-wide association study with the mixed linear model (MLM) using the 3000 phenotypes and genomes of an F1. The individuals of the F1 were generated with random mating of all 1000 individuals of F_0_ => *MLM-3000-F1*
f. Marker-assisted selection using estimated allelic effects of the 100 SNPs with the lowest p-values in a genome-wide association study with the general linear model (GLM) using 3000 unrelated individuals of the founder population=> *GLM-3000-Fd*
g. Marker-assisted selection using estimated allelic effects of the 100 SNPs with the lowest p-values in a genome-wide association (GLM) study using 10,000 unrelated individuals of the founder population=> *GLM-10000-Fd*

All scenarios were repeated 10 times.

## Results

### Genetic gain

The selection based on progeny tests performed best in all tested scenarios, irrespective of the number of causal variants (20 vs. 200) or the level of kinship (0 vs. 0.04). The progeny tests delivered high genetic gains of more than 20% in the first breeding cycle and cumulative genetic gains between 40% and 70% at the end of five cycles (Figure 1 a-c). In both scenarios with a burn-in phase and a resulting kinship structure, genomic selection (gBLUP-singleG and gBLUP-crossG) outperformed the selection by phenotypes. As expected, the single generation genomic selection (gBLUP-singleG) delivered slightly better results compared to the cross generation genomic selection (gBLUP-crossG). Overall, MAS based on different GWAS analyses resulted in the lowest genetic gains. Irrespective of the genetic architecture, MAS could not compete even against simple selection by phenotypes and was clearly outperformed by genomic selection. The only exception where MAS achieved higher genetic gains than genomic selection was the first breeding cycle in a population of unrelated individuals, however, here the selection based on phenotypes was on a similar level. When the target trait is controlled by a small number of causal variants (such as 20 SNPs in scenario 2) MAS can lead to rapid fixation of the identified alleles in the breeding population and prevent further genetic gains. This is illustrated by using allelic effects estimated by a GWAS with 3,000 individuals in an F1 (MLM-3000-F1) where no additional genetic gain was realized after breeding cycle 1. It should be taken into account, however, that selection based on GWAS in the founder populations before the burn-in phase (GLM-3000-Fd and GLM-10000-Fd) does not involve any additional phenotyping or genotyping during the breeding cycles. This can be a crucial advantage when selected seed is harvested in heterogenous forest stands where reliable phenotypes cannot be easily assessed due to high levels of environmental variation.

**Figure 1:**
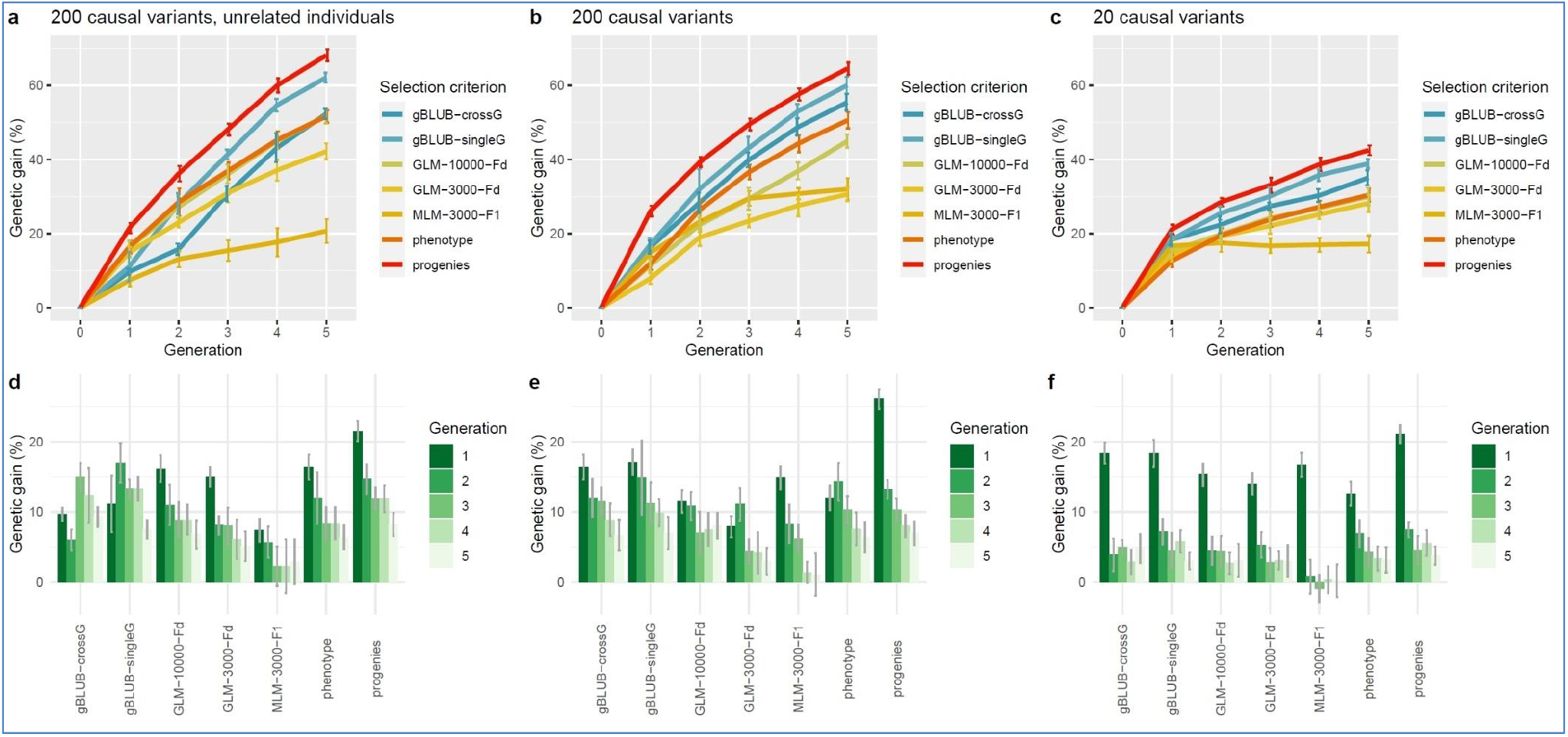
Genetic gains achieved by different selection criteria over five generations in three simulated breeding populations. (**a**, **b**, **c**) Average cumulative genetic gains (n=10) over five breeding cycles are shown for selection based on gBLUP, i.e. genomic selection (blue), GWAS, i.e. marker-assisted selection (green and yellow), phenotypes (orange) and progeny tests (red). (**d**, **e**, **f**) Average genetic gains (n=10) per breeding cycle (Generation) are shown for the different selection criteria. Error bars represent the standard deviation of the mean. The three simulated breeding populations exhibit different genetic architectures, with 200 (a, b, d, e) vs. 20 (c, f) causal variants, and different levels of kinship, with unrelated individuals (a, d) vs. average kinship values of approximately 0.04 (b, c, e, f).

### Precision of GWAS

In each scenario we used MAS with allelic effects estimated in three different GWAS sub-scenarios. First, we ran a mixed linear model (MLM) using the genomic data and kinship information for 3,000 individuals created in an F1 (MLM-3000-F1). In addition, we estimated allelic effects using two general linear model (GLM) analyses with 3,000 (GLM-3000-Fd) and 10,000 (GLM-10000-Fd) unrelated individuals from the founder population before the twenty generations of burn-in. In order to estimate the potential for a GWAS using extreme phenotypes we selected from 10,000 simulated phenotypes the largest 1,500 and smallest 1,500 and repeated the GLM (GLM-3000-extreme). The probability thresholds were set so that for each of the nine GWAS the top 100 SNPs were selected (table 1). Generally, the use of unrelated individuals in the GWAS outperformed the use of related individuals in terms of proportion of true positive SNPs and cumulative genetic gain after the five breeding cycles (table 1). For the scenarios with 200 causal SNPS the increase of the number of individuals in the GWAS led to a higher number of identified SNPs, more precise estimates of the allelic effects and higher genetic gains. Interestingly, nearly the same proportions of identified true positive SNPs and genetic gains were realised with a GWAS using only the 3,000 extreme phenotypes (GLM-3000-extreme). The correlation between the estimated and true allelic effects varied between 0.60 and 0.96 and exceeded 0.9 in all sub-scenarios using 10,000 individuals or extreme phenotypes (“r allele effects” in table 1).

**Table 1:** Results of a post-ex analysis of the number and proportion of true positive SNPs (N true SNPs, % true SNPs) in the simulated GWAS, the Pearson’s correlation coefficient between estimated and true allelic effects (r allele effects) and mean cumulative genetic gain after five breeding cycles (Cumulative genetic gain F5 (%))

| No | Scenario | Name GWAS | p threshold (-log 10)<br>for top 100 SNPs | N true<br>SNPs | % true<br>SNPs | r allele<br>effects | Cumulative genetic<br>gain F5 (%) |
| --- | --- | --- | --- | --- | --- | --- | --- |
| 1 | 1 | MLM-3000-F1 | 4.0017 | 11 | 5.5 | 0.88 | 20.7 |
| 2 | 2 | MLM-3000-F1 | 4.2420 | 5 | 2.5 | 0.60 | 32.0 |
| 3 | 3 | MLM-3000-F1 | 7.9000 | 4 | 20.0 | 0.96 | 17.2 |
| 4 | 1 | GLM-3000-Fd | 4.3800 | 28 | 14.0 | 0.87 | 42.2 |
| 5 | 2 | GLM-3000-Fd | 4.5410 | 23 | 12.5 | 0.79 | 30.6 |
| 6 | 3 | GLM-3000-Fd | 4.8717 | 12 | 60.0 | 0.84 | 28.2 |
| 7 | 1 | GLM-3000-extreme | 5.7800 | 44 | 22.0 | 0.94 | 50.5 |
| 8 | 2 | GLM-3000-extreme | 6.0700 | 43 | 21.5 | 0.96 | 44.3 |
| 9 | 3 | GLM-3000-extreme | 9.9100 | 11 | 55.0 | 0.96 | 25.8 |
| 10 | 1 | GLM-10000-Fd | 7.1800 | 45 | 22.5 | 0.91 | 51.8 |
| 11 | 2 | GLM-10000-Fd | 7.6800 | 45 | 22.5 | 0.96 | 45.0 |
| 12 | 3 | GLM-10000-Fd | 11.9000 | 11 | 55.0 | 0.95 | 30.0 |

### Inbreeding

In all tested scenarios, the inbreeding increased in each breeding cycle (figure 2 a-c). However, the increase was different depending on the selection criteria. In the scenario 1 with 200 causal variants and no burn-in, inbreeding values between 0.051 and 0.117 were observed in the fifth breeding cycle with average Δ*F* per breeding cycle of 0.010 to 0.023 (figure 2a). In scenario 2 and 3 with 200 and 20 causal variants and a burn-in, inbreeding reached values between 0.072 and 0.122 (Δ*F* per breeding cycle of 0.007 to 0.017) in scenario 2 and values between 0.065 and 0.117 in scenario 3 (Δ*F* per breeding cycle of 0.006 to 0.017). The large genetic gains of the genomic selection are linked with a stronger increase of inbreeding (figure 3). The selection based on progeny tests had the best combination of high genetic gains and low levels of inbreeding. The selection based on GWAS resulted in lower levels of inbreeding compared to genomic selection but was in most cases not as effective in terms of genetic gains, as detailed above.

**Figure 2:**
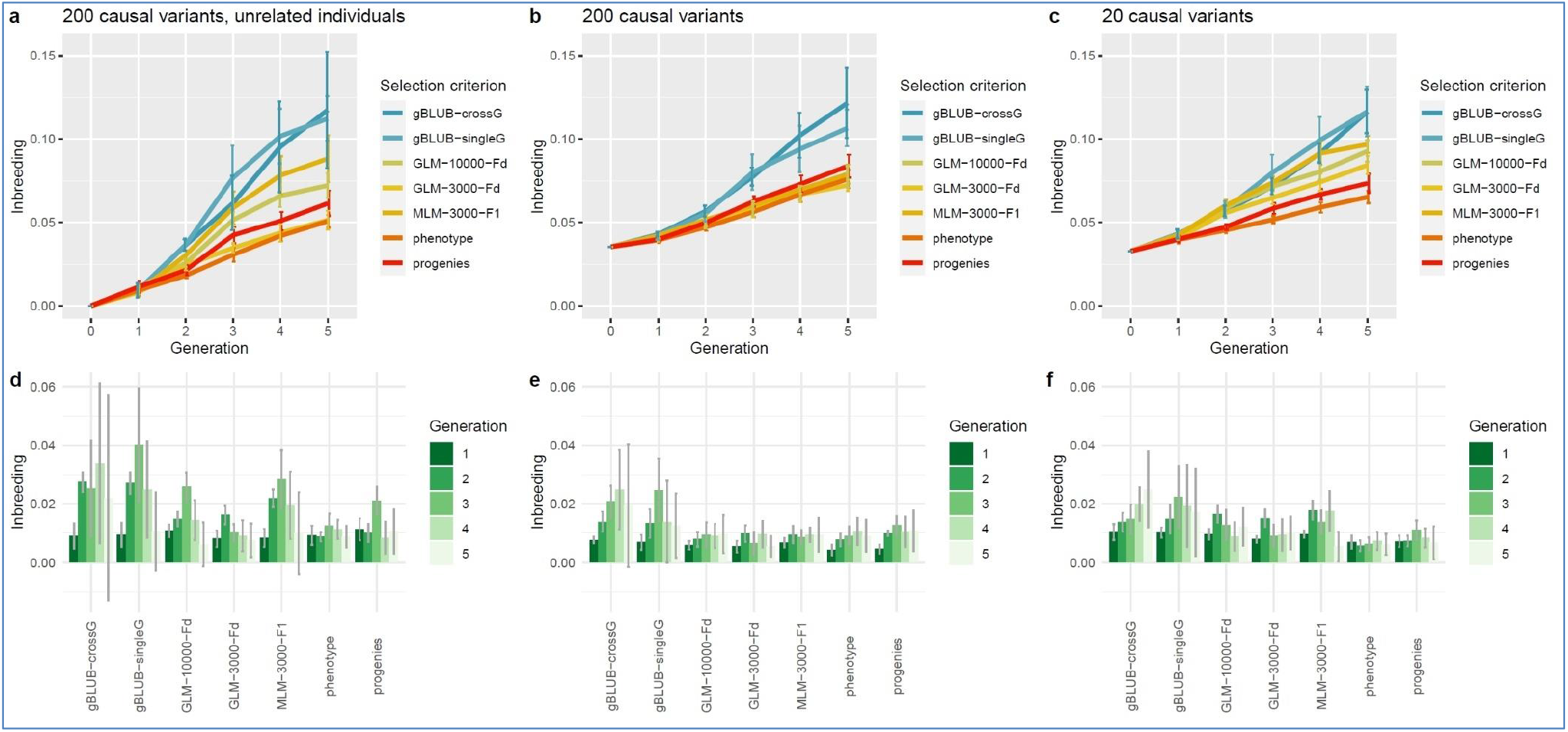
Inbreeding dynamics under different selection criteria over five generations in three simulated breeding populations. (**a**, **b**, **c**) Average cumulative inbreeding values (n=10) over five breeding cycles are shown for selection based on gBLUP, i.e. genomic selection (blue), GWAS, i.e. marker-assisted selection (green and yellow), phenotypes (orange) and progeny tests (red). (**d**, **e**, **f**) Average inbreeding values (n=10) per breeding cycle (Generation) are shown for the different selection criteria. Error bars represent the standard deviation of the mean. The three simulated breeding populations exhibit different genetic architectures, with 200 (a, b, d, e) vs. 20 (c, f) causal variants, and different levels of kinship, with unrelated individuals (a, d) vs. average kinship values of approximately 0.04 (b, c, e, f).

**Figure 3:**
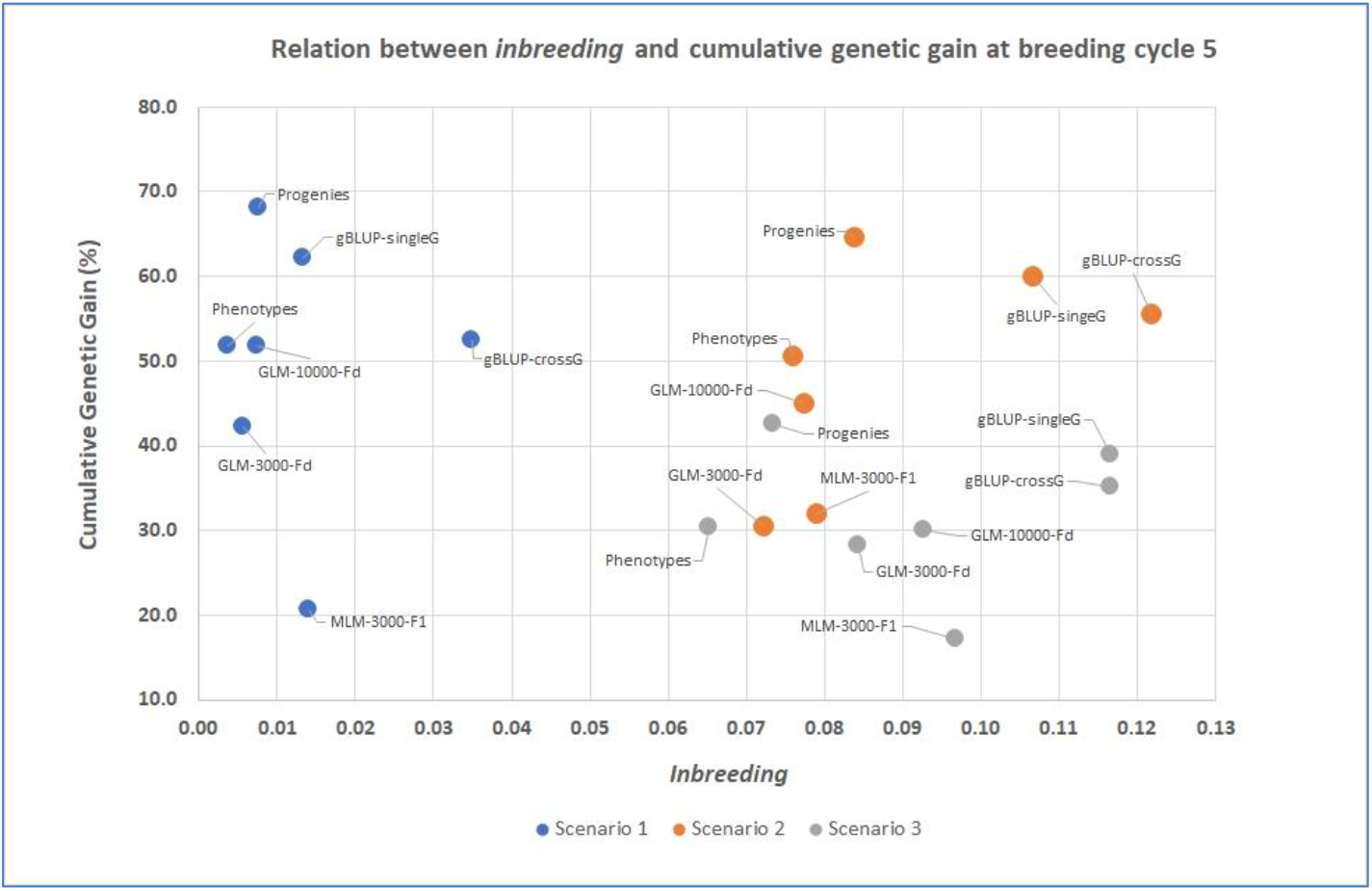
Relation between inbreeding and the cumulative genetic gain (%) at the end of breeding cycle 5 in three different breeding scenarios. Scenario 1 (blue) with 200 causal SNPs and no burn in, scenario 2 (orange) with 200 causal SNPs and burn in and scenario 3 (grey) with 20 causal SNPs and burn in.

## Discussion

### What makes SNPscan special?

Although there are quite a few other simulation programs and R-packages available to simulate breeding strategies and to study sample strategies, SNPscan has special features that make it a useful tool. For example, it is a user-friendly windows application that allows unexperienced users to get started quickly. Possible disadvantages of a windows application in terms of computing speed are compensated by the broad use of parallel programming that enables a maximum number of CPU cores at the same time. The main focus of the program is on the forward simulations with many different criteria to select the parents used for the next breeding cycle. To this end, selection by phenotypes, breeding values, genomic selection and MAS based on allelic effects estimated by GWAS are implemented. Import and export options are available to directly interact with the widely used software TASSEL (Bradbury et al. 2007) and the R-package rrBLUP (Endelman 2011). A unique feature is the post-ex analysis of GWAS performance and the conclusions that can be drawn on the effectiveness of a given sample strategy and used GWAS algorithm to identify SNPs and estimate allelic effects for user defined genetic architectures.

Compared to other software such as GeneEvolve (Tahmasbi and Keller 2017) SNPscan makes strong simplifications of the genome but still maintains important characteristics such as total genome size, number of SNPs, segregation and crossing-over that allow the study of different sample strategies. The software XSim version 2 (Chen et al. 2022) and ADAM-Plant (Liu et al. 2019) are more sophisticated with regard to the simulation of genomic selection in a breeding program and complex crossing schemes but they do not implement MAS based on actual GWAS results.

### Good performance of genomic selection and moderate gains from MAS based on GWAS

In our simulation study the genomic selection performed better in almost all cases compared to MAS scenarios using allelic effects estimated with GWAS. The cross generation genomic selection (gBLUP crossG) provided nearly the same genetic gains as the single generation genomic selection (gBLUP singleG) in most cases. This is of high relevance for practical breeding programs with trees because it suggests that an effective selection of future parents can be done without phenotypes and thus several years earlier. The MAS based on GWAS with large sample sizes of unrelated individuals outperformed genomic selection (first breeding cycles in scenario 1) only when there was no or only a weak kinship structure. Our results on GWAS using extreme phenotypes indicate that a strong reduction of sample size is possible without losing much performance.

Our findings are in accordance with many other publications showing the usefulness of genomic selection in breeding programs. This has been demonstrated with simulation studies (Meuwissen et al. 2001, Grattapaglia and Resende 2011) and successfully implemented in practical breeding operations first in dairy cattle breeding programs (Su et al. 2010), then in agricultural crops (Robertsen et al. 2019) and finally in forestry species (El-Kassaby et al. 2012, Grattapaglia et al. 2018).

So far, the application of marker-assisted selection in forest breeding programs could not be successfully established. Grattapaglia et al. (2018) stated that while marker-assisted selection was mostly fruitless, genomic selection has been proven to be very successful. More positive conclusions on the application of GWAS results in breeding have been drawn from crop (Cortes et al. 2021, Saini et al. 2022) and animal breeding (Gutierrez-Reinoso et al. 2021). The main explanation for the difficulties in the application of marker-assisted selection has been the undiscovered complexity of the genetic architectures of the target traits with many causal loci with small effects and low frequency. SNPs with larger effects and SNPs with moderate effects but higher allele frequencies are picked up in our simulations by GWAS. These alleles become fixed in the first breeding cycles and thereby limit additional genetic gains in the following cycles. With high numbers of unrelated individuals (> 10,000) more causal SNPs can be identified but because of their small allelic effects and low frequencies their impact on the genetic gain is small. In most practical breeding programs phenotyping is much cheaper than whole genome sequencing. We did some first promising simulations on GWAS with extreme phenotypes (e.G. the 15% edges of the phenotypic distribution). Here we observe higher genetic gains even with relatively small sample sizes of 3,000 individuals, however, these scenarios need to be studied in more detail.

One important aspect to consider when interpreting our simulation results and assessing possible breeding strategies is the phenotyping. Both, the genomic selection and the phenotypic selection rely on repeated phenotyping of the breeding population. In our simulations using MAS, we computed a GWAS only once for a diverse founder population of unrelated individuals. The results from this single GWAS were then used during the entire simulated breeding program. Depending on the target species, this methodological difference can have a profound impact on the feasibility of the whole program. Thus, the repeated phenotyping needed in genomic selection to keep the training population accurate will delay the breeding program by several years compared to selection by MAS. Moreover, there is the advantage of MAS to be more suitable for the integration of new unrelated material into a breeding program and whenever phenotyping is limiting.

#### Critical increase of inbreeding

In the simulated breeding program, the increase of inbreeding per breeding cycle (Δ*F*) varied in the different scenarios between 0.006 and 0.023. The highest Δ*F*-values were observed for the scenarios with genomic selection. The optimization of a maximum genetic gain and minimal inbreeding and thus low loss of genetic diversity has been a challenge for most breeding programs. Generally, changes of the selected individuals for the mating and particular crossing schemes using mathematical algorithm are applied to achieve an optimization (Woolliams et al. 2015). Usually breeding programs, especially animal breeding programs, mitigate inbreeding by optimum contribution selection (OCS). This methods aims to keep the average coancestry of the selected parents at a certain level and thus controls the short term and long term inbreeding (Meuwissen 1997). The OCS approach has been improved in several steps and the most recent methods also consider different levels of genetic introgression (Kohl et al. 2020).

Our simulated Δ*F*-values are similar to real breeding programs with comparable census numbers and selection intensity in the breeding population. For example, the breeding program of the tree species *Pinus taeda* in North Carolina started with 935 selected trees and controlled the inbreeding not to exceed 0.0625 (Isik and McKeand 2019). In animal breeding the negative effects of genomic selection on inbreeding have been shown with simulation studies as well (Sonesson et al. 2012). In a detailed study on the negative impact of inbreeding Wu et al. (2016) simulated seven different tree breeding strategies. They included inbreeding depression caused by deleterious alleles in their study. They concluded that forced inbreeding will more likely lead to fixation of deleterious alleles and to a lesser extent to a pruning effect of those negative alleles. The optimal balance between the benefits of increased genetic gain and the drawbacks of elevated levels of inbreeding needs to be carefully considered for each specific breeding program.

## Conclusions

Using simulations of different breeding populations and strategies, our results further support the potential of genomic selection for forest tree breeding and improvement. Nevertheless, using whole-genome data of large sample sizes or extreme phenotypes for GWAS may provide advantages over genomic selection under certain conditions and could revive efforts for marker-assisted selection, especially when phenotyping represents a bottleneck. We will focus future work on the integration of real genome data with SNPscan to enable the simulation of more realistic breeding programs such as breeding programs with infusion of unrelated individuals and the sub-division in several breeding populations. For further testing the potential of MAS we will create additional links to other GWAS analysis programs such as GAPIT (Wang and Zhang 2021) for more advanced GWAS algorithm such as BLINK (Huang et al. 2019). Additionally, we will study in more detail the possibilities of MAS based on GWAS with extreme phenotypes. Considering the broad implementation of genomic selection methods in the field of forest tree breeding in the last decade, it will be exciting to follow the impacts on the actual forest ecosystems and further develop strategies to adapt forest tree species to the rapidly changing environmental conditions. In forestry the genomic revolution has only just begun.

## Data availability

SnpScan has been programmed with Visual Studio 2019 as a.NET application (framework 4.7.2) and compiled as 64-bit versions for the operating system Microsoft Windows (Windows 10). The program, the user manual and different videos that explain the program are available on our website: https://www.thuenen.de/en/institutes/forest-genetics/software/SNPscan

## Acknowledgement

We are thankful to members of the Centre for Integrated Breeding Research (CiBreed) at the University of Göttingen for helpful discussions on SNPScan. We would like to thank Malte Mader for critical testing of the program and for helpful suggestions of its improvement.

